# Clinically observed deletions in SARS-CoV-2 Nsp1 affect protein stability and its ability to inhibit translation

**DOI:** 10.1101/2021.11.03.467065

**Authors:** Pravin Kumar, Erin Schexnaydre, Karim Rafie, Ilya Terenin, Vasili Hauryliuk, Lars-Anders Carlson

## Abstract

Nonstructural protein 1 (Nsp1) is a major pathogenicity factor of SARS-CoV-2. It inhibits host-cell translation, primarily through a direct interaction between its C-terminal domain and the mRNA entry channel of the 40S small ribosomal subunit, with an N-terminal β-barrel domain fine-tuning the inhibition and promoting selective translation of viral mRNA. SARS-CoV-2 *nsp1* is a target of recurring deletions, some of which are associated with altered COVID-19 disease progression. To provide the biochemical basis for this, it is essential to characterize the efficiency of translational inhibition by the said protein variants. Here, we use an *in vitro* translation system to investigate the translation inhibition capacity of a series of clinically observed Nsp1 deletion variants. We find that a frequently observed deletion of residues 79-89 destabilized the N-terminal domain (NTD) and severely reduced the capacity of Nsp1 to inhibit translation. Interestingly, shorter deletions in the same region have been reported to effect the type I interferon response but did not affect translation inhibition, indicating a possible translation-independent role of the Nsp1 NTD in interferon response modulation. Taken together, our data provide a mechanistic basis for understanding how deletions in Nsp1 influence SARS-CoV-2 induction of interferon response and COVID-19 progression.

## Background

Coronaviruses (CoVs) are a family of enveloped, positive-sense single-stranded RNA viruses subdivided into four genera: Alpha-, Beta-, Gamma- and Deltacoronaviruses (1). Four so-called common-cold cornaviruses that cause only mild disease are prevalently circulating in the human population: HCoV-OC43, HCoV-HKU1, HCoV-229E and HCoV-NL63 (2). The situation has changed dramatically in the recent years with the zoonotic introduction of three highly pathogenic Betacoronaviruses into humans: SARS-CoV in 2002 (2), MERS-CoV in 2012 (2) and – most recently – SARS-CoV-2 in 2019 (3–6). SARS-CoV-2 is the cause of the COVID-19 pandemic which to date has caused nearly five million confirmed deaths (https://coronavirus.jhu.edu/map.html, retrieved 2021-10-21).

Coronaviruses have unusually large genomes for positive-sense single-stranded RNA viruses. At 27-32 kb they are an approximately threefold larger than other representatives of this group. This genomic expansion is thought to be made possible by a more complex, proofreading-capable polymerase (7), and it has allowed coronaviruses to acquire a larger toolbox of host-cell-manipulating proteins. One such key pathogenicity factor is the nonstructural protein 1 (Nsp1). This protein has no known enzymatic activity, and while betacoronaviruses deleted for the *nsp1* gene can replicate in cell culture, they are strongly attenuated *in vivo* (8). The 180 amino acid-long SARS-CoV Nsp1 was found to suppress the antiviral host-cell interferon response through a dual mechanism: it mediates the cleavage of host-cell mRNAs by an unknown ribonuclease (9), and it suppresses translation through direct interaction with the large (40S) subunit of the ribosome (10). Distinct mutations in *nsp1* have been described that selectively abrogate either of these two effects: substitutions of Arg124-Lys125 in the folded N-terminal domain of SARS-CoV were shown to abolish the Nsp1-induced mRNA cleavage (11), and residues Lys164-His165 in the C-terminal domain have been shown to be essential for binding to the 40S subunit of the ribosome (Fig. 1A) (12,13). SARS-CoV-2 Nsp1 binds empty small ribosomal subunit, as well as the 43S preinitiation complex and the full 80S ribosome, in all cases only efficiently engaging the open conformation of the small subunit (14). In cryo-EM structures of SARS-CoV-2 Nsp1 bound to the human 40S, the C-terminal ~30 residues (148-180 in PDB ID 6ZLW) were present in a sufficiently rigid conformation that an atomic model could be built (15–17). This model shows that the C-terminal part of Nsp1 folds as two α-helices that form a hairpin-like arrangement inside the mRNA tunnel, incompatible with concomitant binding of mRNA. Consistent with the functionally crucial location of the C-terminal domain in the Nsp1:40S complex, both its truncation (Δ118-180) and K164/H165A substitution abrogate the Nsp1 interaction with the small ribosomal subunit (18). Off the ribosome the C-terminal region (131-180) is unstructured, suggesting a possibility that the structure is attained upon recruitment to the ribosome (19). The N-terminal domain could be located in the cryo-EM maps but at too low a resolution to allow model building, presumably due to flexibility with respect to the 40S. The cryo-EM derived location of the Nsp1 N-terminal domain in the ribosomal complex is further supported by *in situ* cross-linking mass spectrometry identifying multiple crosslinks between Nsp1 and ribosomal protein S3 (18,20).

**Figure 1:**
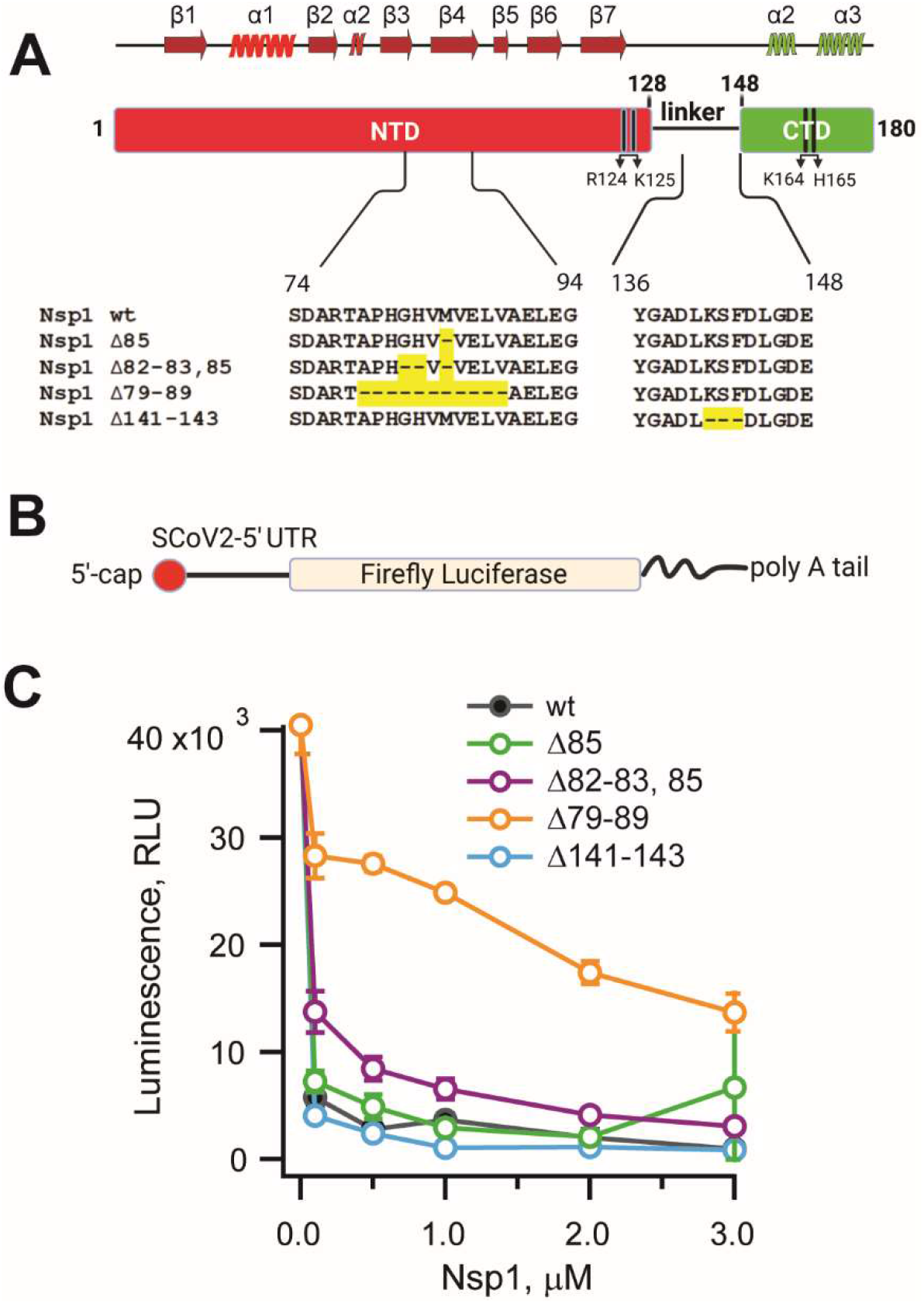
Nsp1 variants with amino acid deletions located outside of the mRNA tunnel-targeting CTD differ in their ability to inhibit translation. **(A)** Structural organization of SARS-CoV-2 Nsp1 protein showing the N-terminal and C-terminal domains (NTD and CTD) as well as the regions containing naturally occurring deletions (25,26). Also denoted are amino acids R124-K125 necessary for host mRNA cleavage (11) and the R164-K165 pair reported to be necessary for the interaction with the 40S mRNA entry site (12). The secondary structure elements of wild-type Nsp1 are shown above the domain organization. **(B)** Schematic of the mRNA reporter construct containing the 5’ UTR of SARS-CoV-2 followed by a firefly luciferase open reading frame, with 5’ cap and 3’ poly A tail. **(C)** Results of the *in vitro* translation assay to assess the translation efficiency of the reporter mRNA. Graph showing luminescence signal in response to firefly luciferase activity in HEK293F translational lysate at the end of the reaction, in the absence and presence of increasing concentrations (0.1, 0.5, 1.0, 2.0, and 3.0 μM) of recombinant Nsp1 and deletion variants (Δ85; Δ82-83,85; Δ79-89; Δ141-143). The average luminescence (RLU) from three independent experiments are shown in the plot, error bars represent standard deviation.

The structure of Nsp1 bound to the ribosome raises the question how any translation is possible if Nsp1 blocks the mRNA channel? While the answer to this question still appears incomplete, some important findings have been made. First, based on *in vitro* translation systems SARS-CoV-2 Nsp1 appears to inhibit translation of both host and SARS-CoV-2 mRNAs, but mRNAs with the viral 5’ UTR less so (14,17,21). Second, the mRNAs of SARS-CoV-2 and related SARS-CoV appear to partially escape Nsp1 suppression through an interaction between Nsp1 and the stem-loop region 1 (SL1) of the viral 5’ untranslated region (UTR) (21,22). The interaction is suggested to only take place on the 40S subunit since isolated Nsp1 has not affinity for the 5’ UTR (21). For SARS-CoV Nsp1 it was shown that the Arg124-Lys125 residues, which are necessary for Nsp1-induced cleavage of cellular mRNAs, are equally necessary for the 5’ UTR mediated escape of Nsp1 suppression (22). The N-terminal region plays an important role in both shutting off host translation as well as in sustaining robust 5’ UTR-dependent expression of the viral mRNA, with substitutions (R124A/K125A, E36A/E37A) and deletion (Δ1-117) in N-terminal domain attenuating both the ribosome and mRNA binding (18).

In the course of the ongoing COVID-19 pandemic, extensive genetic characterization has been carried out of new emerging mutants and virus variants. Nsp1 displays one of the highest degrees of diversity amongst SARS-CoV-2 proteins (23–25), with several circulating viruses with deletions in Nsp1 reported (25,26). A three-residue deletion (Δ141-143) was described as appearing in clinical isolates from different geographical localities (26). These three residues are located close to, albeit not in, the region which forms an ordered structure inside the 40S mRNA tunnel. It thus seems possible that this shortening of the linker region between the ribosome-inserting part and the stably folded N-terminal β-barrel domain (27,28) may affect the association of this altered protein with the ribosome, and thus its effectiveness in translational shutoff. Another cluster of deletions is located around residues 79-89 of Nsp1 (25). This results in various length deletions around a loop in the atypical β-barrel of the folded N-terminal domain. The longest deletion, Δ79-89, is reported to be the fifth most common deletion in SARS-CoV-2 Nsp1 in a worldwide comparison (25). The 79-89 deletion, as well as shorter deletions in the same region of Nsp1, correlate with higher cycle threshold (Ct) values in patients (i.e. lower viral load), less severe disease outcome, and a weaker interferon response as measured by lower serum levels of IFN-β (25). The biochemical basis for the altered disease course associated with Nsp1 deletions is not established. Here, we purified a number of Nsp1 proteins corresponding to these circulating mutations in SARS-CoV-2, and compared their potency in inhibiting translations using a human cell *in vitro* translation lysate. We correlate the findings to thermal stability assays and protein fold predictions of the mutants. The result shed light on the clinical and cellular findings related to Nsp1-mutated SARS-CoV-2 isolates.

## Results

### Circulating deletions in SARS-CoV-2 Nsp1 differ in their inhibitory effect on translation

Several amino acid deletions are reported in SARS-CoV-2 Nsp1: Δ85, Δ82-83, 85 and Δ79-89 in the N-terminal domain (NTD) (25) and Δ141-143 in the C-terminal domain (CTD) (26) (Fig. 1A). These deletions have been correlated to clinical characteristics, but the mechanistic biochemical basis of why they alter the course of COVID-19 has not been determined. We wished to assess the effect of these deletions on Nsp1’s ability to shut down host translation. As a first step towards this analysis, we purified wild-type Nsp1 and a series of deletion mutants to homogeneity and monodispersity (Fig. S1). We then established a translation assay based on lysates of the human cell line HEK293F. Since Nsp1 has been reported to reduce translation of viral mRNAs to a lesser degree than cellular mRNAs (14,17,21), we reasoned that a reporter mRNA resembling a viral mRNA would provide a more sensitive assay. Thus, we measured translation efficiency of a reporter mRNA encoding firefly luciferase, equipped with the SARS-CoV-2 5’ UTR, a 5’ cap, and 3’ poly(A) sequence, in the presence of increasing concentrations of both wild-type recombinant Nsp1 and its deletion variants (Fig.1B).

In agreement with earlier reports (17,21), wild-type Nsp1 efficiently abrogates production of firefly luciferase in concentration-dependent manner. Already at 0.1 μM, wild-type (wt) protein reduces the translational efficiency by more than 95% (Fig. 1C). An effect similar to the wt is observed for the NTD deletion variant Δ85 and the CTD deletion variant Δ141-143. The NTD deletion variant Δ82-83,85 shows slightly reduced suppression of translation compared to the wild type. In contrast, the longest deletion in the NTD, Δ79-89, is substantially weakened in its ability to inhibit translation as compared to wt Nsp1 (Fig. 1C). We observed that even at 3 μM protein concentration, the translation efficiency reduces only to 40-50 %, establishing that the Δ79-89 protein is less effective in translation shutdown than the wild-type and other deletion variants. Taken together, these data establish a biochemical basis for correlating COVID-19 disease progression to the translational shutdown efficiency of circulating deletion mutants in Nsp1.

### Deletions in Nsp1 lead to altered protein stability that correlates with translation inhibition

None of the investigated *nsp1* mutations was in the region previously reported to interact with the ribosome, and yet the longer deletions in the NTD were clearly altered in their translation shutdown capacity. We wanted to investigate if this effect could be due to the destabilizing effects of the deletions on the NTD. We first noted that all proteins were soluble and monodisperse when purified from *E. coli*, but the Δ79-89 construct eluted at a lower volume in size exclusion chromatography, indicting a larger hydrodynamic radius (Fig. S1). To probe the effects of the deletions on protein stability, we assessed the thermal stability of wt Nsp1 and deletion variants by thermal shift assay (TSA) (29) (Fig. 2). In this assay, SYPRO orange dye is used as a probe to estimate the extent of unfolding of the protein with increasing temperature, and the melting temperature (T_m_) from each curve is determined as described in the experimental procedure section. Raw TSA data (Fig. 2A, B) demonstrates that all mutants of SARS-CoV-2 Nsp1 showed similar, wild-type-like unfolding profile, apart from the longest deletion variant (Δ79-89). Wt Nsp1 and all altered proteins, with a notable exception of Δ79-89, allowed for fitting of a simple melting curve (Fig. 2C) indicating a single transition temperature that is comparable to that of the wt protein (Fig. 2E). The thermal denaturation curve of Δ79-89 (Fig. 2A,B) was shallow and could not be perfectly recapitulated by fit to a single melting temperature (Fig. 2D). However, using the first part of the curve gave a fit that is consistent with the qualitative appearance of the curve (Fig. 2D) and a lowered thermal stability. This suggests that already at room temperature the structural integrity of Δ79-89 variant is compromised. In summary, our results establish a correlation between structural stability of Nsp1’s NTD and its ability to inhibit translation.

**Figure 2.**
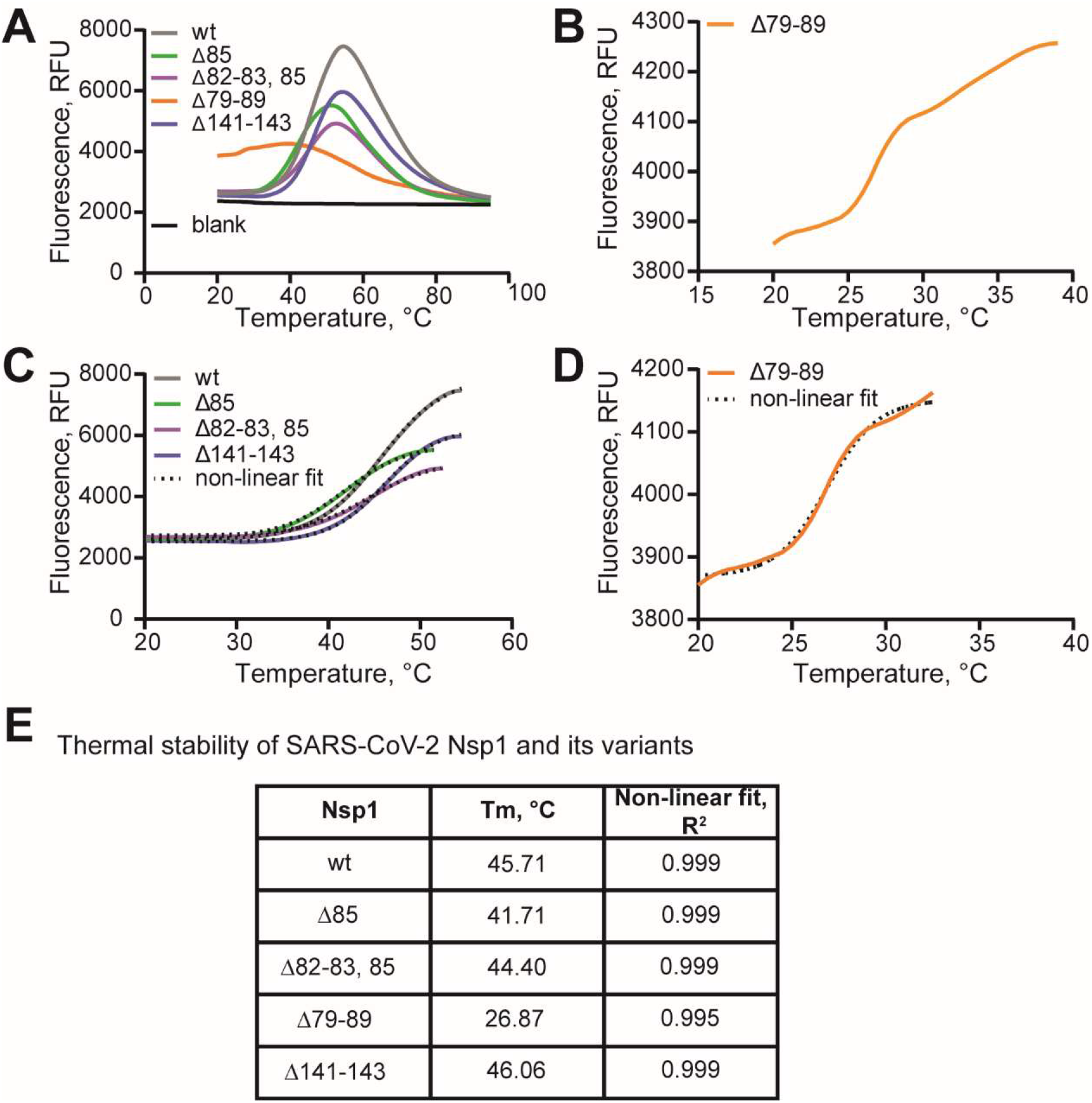
The Δ79-89 deletion variant of SARS-CoV2 Nsp1 has a drastically reduced thermal stability. **(A)** Raw data of thermal shift assays (35) with Nsp1 variants using SYPRO Orange. **(B)** Raw TSA data of the deletion variant Δ79-89. **(C)** Non-linear regression curve fit (Boltzman sigmoidal) of the unfolding curves of Nsp1 variants (except that of the Δ79-89 protein) to calculate the melting temperature. **(D)** Same as (C), for the Δ79-89 variant. **(E)** Tabular presentation of the result of the non-linear fitting of the unfolding curve of all the proteins.

### Structure predictions suggest that NTD β-barrel destabilization causes the decreased stability of Δ79-89 nsp1

We next wanted to explore the possible structural basis for the decreased translation inhibition and thermal stability of the altered Nsp1 proteins. To investigate how the Nsp1 deletions impact its structure we predicted the structures of wt, Δ141-143, Δ85, Δ82-83, 85 and Δ79-89 Nsp1 using AlphaFold 2 (30). Since there are available experimental structure of the free SARS-CoV-2 Nsp1 NTD (residues 10-126) (27), as well as the ribosome-inserted CTD (15–17), we could compare them to the prediction of the wt Nsp1 structure as a baseline. The predicted Nsp1 structure aligns very well with the experimentally determined N-terminal β-barrel domain, with average positional shifts (root-mean-square deviation, RMSD) of the respective Cα atoms of only 0.64 Å for residues 10 to 126 (Fig. 3A, Fig. S2A). In the CTD, two α-helices are correctly predicted where they are observed in the experimental structure, albeit at a different angle to each other than in the 40S-bound structures (Fig. S2A). Overall, the per-residue prediction quality score (pLDDT) correlated with the rigidity of the fold across Nsp1, with the NTD β-barrel having higher scores than the CTD which was reported to be flexible in solution (Fig. S2B) (19). Supported by the accurate prediction of the wt Nsp1 structure, we thus reasoned that structure predictions for the altered Nsp1 proteins may shed light on the differential effects of the deletions on protein stability. Unsurprisingly, the deletion Δ141-143, predicted to be located in a disordered region of the CTD, had no effect on the predicted fold of other parts of Nsp1 (Fig. S2). The other three deletions (Δ85, Δ82-83, 85 and Δ79-89) are all located at the beginning of the fourth β-strand of the β-barrel (β4) (Fig. 3B, D). Interestingly, despite being located in a secondary structure element, the two shorter deletions are not predicted to affect the β-barrel integrity (Fig. S2C), likely given that the hydrogen bonding between β-strands is mediated by the protein backbone and can be rescued by the residues “next-in-line” to the deleted residues. The predicted integrity of the fold of these deletions correlates well with their unaltered thermal stability (Fig. 2E). The only striking effect on the integrity of the β-barrel domain is predicted to stem from the longest deletion. The Δ79-89 structure prediction suggests a near-complete dissolution of the β3-β5 strands leading to a break in the β-barrel domain, severely affecting its integrity (Fig. 3C, E). This prediction is in line with the drastic reduction in thermal stability for the Δ79-89 protein from 46°C to ~27°C (Fig. 2E). Taken together, structure predictions of the Nsp1 deletions suggest destabilization of the N-terminal β-barrel domain as the mechanism behind the altered properties of the Δ79-89 Nsp1 protein.

**Fig. 3:**
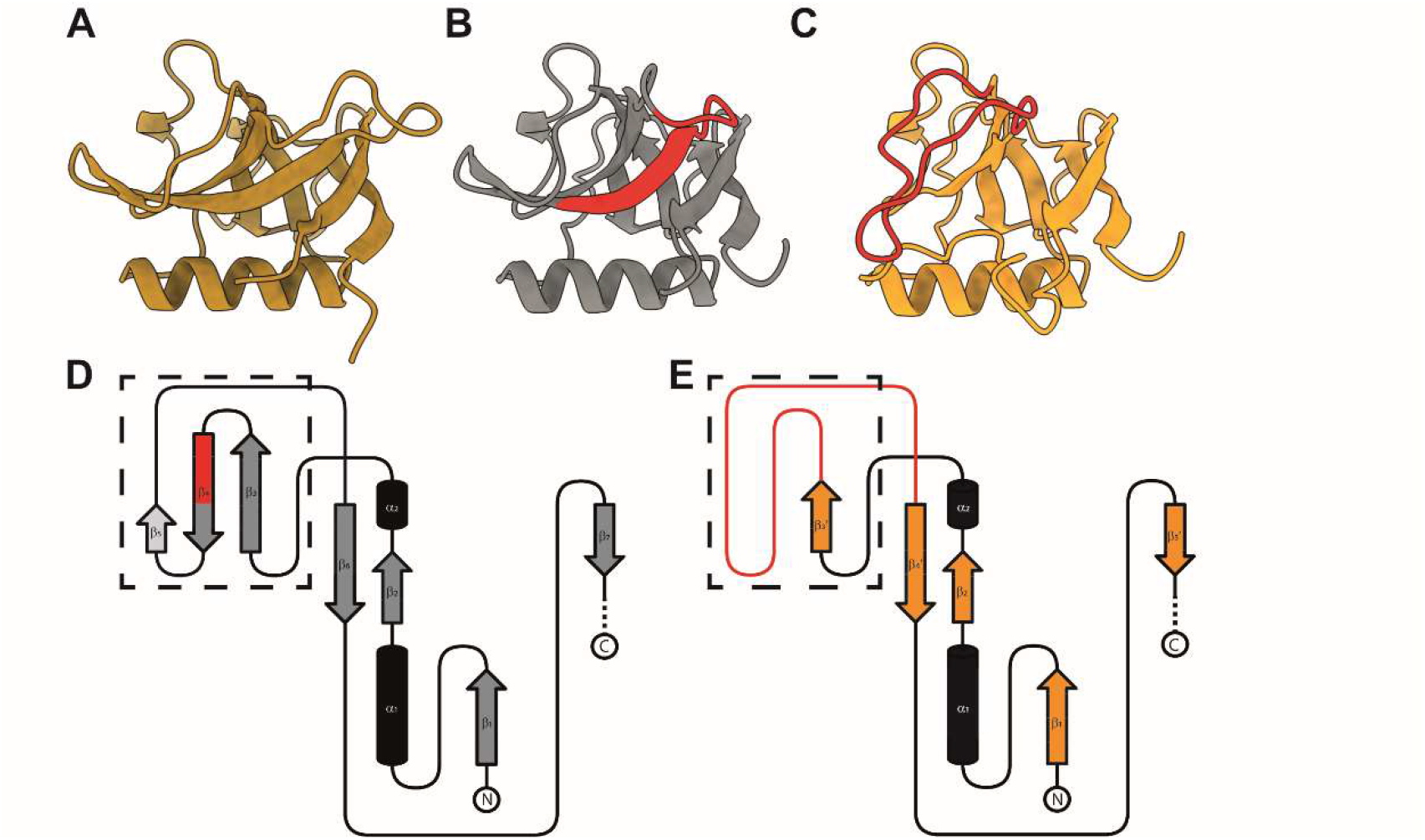
Predicted β-barrel integrity correlates with altered thermal stability of Nsp1 deletion variants. **(A)** Cartoon representation of the experimentally determined structure of the Nsp1 N-terminal β-barrel domain (PDBID: 7K7P, (27)). **(B)** Cartoon representation of the predicted wt Nsp1 structure (grey), truncated to the same boundaries as for (A). In red highlighted are residues 79-89. **(C)** Cartoon representation of the predicted Δ79-89 Nsp1 structure (orange), truncated to the same boundaries as for (A). In red highlighted is the disordered region, which in (B) forms the β-strands β3-β5. **(D)** Topological drawing of the wild-type Nsp1 fold with α-helices shown as black tubes and the β-strands as arrows. β-strands forming the β-barrel are shown in grey and non-β-barrel forming strands in light grey. Highlighted in red is the β-strand part deleted in Δ79-89 Nsp1. **(E)** Topological drawing of the Δ79-89 Nsp1 fold. α-helices are shown as black tubes and the β-strands as orange arrows. Highlighted in red is the disordered region as in (C). **(D-E)** Dashed-lined box highlights the location of change between the predicted wt and Δ79-89 folds.

## Discussion

The high transmissibility of SARS-CoV-2 and the lack of pre-existing immunity contributed to an explosive development of the COVID-19 pandemic. This happening in the age of readily available RNA sequencing has given a fine-grained view of the mutations taking place in the viral genome in the course of the pandemic. The nonstructural protein Nsp1 is a major pathogenicity factor for coronaviruses (8). Although Nsp1 has remained unaltered in the major SARS-CoV-2 variants, various internal deletions in Nsp1 (Fig. 1A) are regularly detected in clinical isolates and have been correlated to different clinical outcomes (25,26). In this study, we used a human translation lysate assay to compare the potency of these Nsp1 deletion mutants in inhibiting translation. Notably, none of the deletions encompassed the two motifs previously identified as being necessary for Nsp1’s ability to inhibit translation (12), and mediate mRNA degradation (11), respectively (Fig. 1A). Thus, it was not clear *a priori* what their effect on translation would be, if any. The main finding of this study is that several of these circulating Nsp1 deletion mutants are still highly potent in inhibiting translation *in vitro*, except for the longest deletion, Δ79-89. Whereas still soluble and monodisperse, the Δ79-89 protein was no longer able to shut down translation completely, even when present at thirtyfold higher concentrations than the wild type protein. We show that Δ79-89 Nsp1 also stands out from the other deletion mutants by having severely reduced thermal stability. Structure predictions of all mutants were consistent with this and suggested that the mechanism for the lost stability of the Δ79-89 protein is a rearrangement leading to a disruption of its N-terminal β-barrel domain fold.

The deletions around residues 79-89 in Nsp1 were previously characterized in terms of their effect on transcriptome and type I interferon response (25). A side-by-side comparison of those results with our translation shutdown data reveals some interesting differences. Strikingly, the shorter deletions in this region already alter the transcriptome and lead to a much weakened type I IFN response as compared to the 180 residue version of Nsp1. These shorter deletions are not impaired in translation shutdown at the concentrations tested, indicating that this region of Nsp1 may have other functions. Notably, the deletions flank the only long loop connecting two strands of the β-barrel. This loop was reported to be flexible in the NMR structure of SARS-CoV Nsp1 (28), but it is to date unknown whether is involved in interactions with host or viral components.

In summary, we determined the translation shutdown capacity of several clinically detected deletions in SARS-CoV-2 Nsp1. The data show that a compromised stability of the N-terminal β-barrel abrogates translational shutdown, and further indicates that the shorter deletions around residue 79-89 may affect interferon response and transcriptome through a mechanism independent of translation shutdown.

### Experimental procedures

#### Cloning and mutagenesis

For expression of wild-type Nsp1, the *nsp1* gene was amplified from the construct pLVX-EF1alpha-SARS-CoV-2-nsp1-2xStrep-IRES-Puro (Addgene) (31) and inserted into a 1B vector (Macrolab, UC Berkeley) using In-Fusion cloning kit (Takara Bio). Deletion mutations were generated from this plasmid by standard site-directed mutagenesis methods. The reporter plasmid T7-5’UTR-Firefly luciferase for RNA synthesis was generated by inserting the T7 promoter upstream of SARS-CoV2 5’ UTR fused to the firefly luciferase coding sequence into the destination vector pUC19. All plasmids were sequenced to confirm cloning of the correct sequence.

#### Protein expression and purification

Wt Nsp1 and all deletion variants were expressed and purified as follows. The plasmid was transformed into *E. coli* BL21(DE3) cells for overexpression. An overnight culture was grown at 37°C to inoculate the secondary culture. Cells were grown at 37°C until the OD_600_ reached 0.4, after which the incubator temperature was changed to 25°C to let the cells cool down to induction temperature 25°C. At OD_600_ of around 0.8-0.9 the protein expression was induced by addition of 0.5 mM Isopropyl β-d-1-thiogalactopyranoside (IPTG) and the protein was expressed at 25°C overnight. Cells were harvested by centrifugation at 6,000 rpm (rotor JLA-8.1000 Beckman Coulter, Brea, USA) for 60 minutes. After discarding the supernatant the cell pellet was washed with lysis buffer (50 mM HEPES-NaOH, pH 7.4, 300 mM NaCl, 0.1 mM THP, 10 mM Imidazole and 5 % glycerol) and stored at −80°C.

Cell mass was thawed and resuspended in lysis buffer supplemented with DNase I and protease inhibitor cocktail (1 mM benzamidine, 0.2 mM phenylmethylsulfonyl fluoride, 5 μM leupeptine). Homogenized suspension was then passed twice through a cell disruptor (Constant System Limited, Daventry, England) at a pressure 27 kPsi. The lysate was clarified by centrifugation at 21,000 rpm (rotor JA-25.50 Beckman Coulter, Brea, USA) for 1 hour, and the supernatant passed through a 0.22 μm syringe filter. 1 ml Ni-Sepharose Fastflow resin (Cytiva) pre-equilibrated with lysis buffer was combined with clarified lysate and incubated for 2 h at 4°C under gentle agitation. Next the resin was loaded onto a gravity-flow column. The protein-bound resin was washed twice with 20 ml wash buffer (50 mM HEPES-NaOH, pH 7.4, 300 mM NaCl, 0.1 mM THP, 30 mM Imidazole and 5% glycerol) twice. The resin was then resuspended in 4 ml lysis buffer, supplemented with TEV protease (approx. 70 μg/ml) and incubated overnight at 4°C on a rotator wheel. The cleaved protein was collected as flowthrough. An additional wash with 5 ml lysis buffer was performed to collect the residual cleaved protein. Both elutions were pooled and further purified by anion exchange chromatography. To do so, after diluting with buffer A (50 mM HEPES-NaOH, pH 7.4, 100 mM NaCl, 0.1 mM THP, and 5% glycerol), diluted sample was filtered using 0.22 μm syringe filter (VWR) and loaded onto a HiTrap Q HP 1ml column (GE healthcare). The sample was eluted with a gradient going from buffer A to buffer B (50 mM HEPES-NaOH, pH 7.4, 1 M NaCl, 0.1 mM THP, and 5% glycerol) in 14 ml. Nsp1-containing fractions were pooled and concentrated using a Vivaspin 6 centrifugal unit with 5 kDa cut off membrane (EMD Millipore) before being loaded onto a Superdex 75 increase 10/300 size-exclusion column (Cytiva) that was pre-equilibrated with SEC buffer (20 mM HEPES-KOH, pH 7.51, 200 mM KOAc, 2 mM Mg(OAc)_2_, 0.1 mM THP, and 2.5 % glycerol). Protein elutions were subjected to SDS-PAGE gel electrophoresis and the Nsp1-containing fractions were pooled and concentrated. Aliquots were then flash frozen in liquid N_2_ and stored at −80°C. The identity of all the proteins was confirmed by trypsin-digestion mass spectrometry.

#### In vitro transcription

Capped and polyadenylated RNA transcripts were synthesized from linearized plasmid (T7–5’-SCoV2-UTR–firefly Luciferase) using the mMESSAGE mMACHINE T7 Ultra kit (Invitrogen) following the manufacturer protocol. Briefly, all the reagents including the 5’cap analog were gently mixed with linearized plasmid and transcription was performed by T7 RNA polymerase at 37°C for 2 hrs. Then, 1 μl of Turbo DNase was gently mixed with the transcription mixture and incubated further at 37°C for 15 min. In order to add poly (A) tail to the 3’ end of the in vitro transcribed RNA, E. coli Poly(A) Polymerase I (E-PAP) was added along with other provided reagents to the transcription mixture and after mixing gently incubated at 37 °C for 45 minutes. According to the manufacturer, this poly(A) tailing adds at least 150 adenines to the 3 ‘end of the mRNA. Finally, the RNA prep was recovered by lithium chloride precipitation and quality-checked by running a sample on a denaturing agarose gel, which resulted in a single band. The capped and poly(A)-tailed RNA was aliquoted in 10 μl volumes and flash frozen in liquid N2 and stored at −80°C.

#### Preparation of HEK293F translation lysate

*In vitro* translation lysates were prepared from HEK293F cells using a previously described protocol (32–34). Cells were scraped and collected by centrifugation for 5 minutes at 600 rpm at 4°C. Cells were washed once with cold PBS (137 mM NaCl, 2.7mM KCl, 100mM Na_2_HPO_4_, 2mM KH_2_PO_4_) and re-suspended in Lysolecithin lysis buffer (20 mM HEPES-KOH, pH 7.4, 100 mM KOAc, 2.2 mM Mg(OAc)_2_, 2 mM DTT, and 0.1 mg/ml lysolecithin), using 1 ml for 8×10^6^ cells. Cells were incubated for 1 min on ice, then immediately centrifuged for 10 sec at 10,000 g at 4°C. The pellet was re-suspended in cold hypotonic extraction buffer (20 mM HEPES-KOH, pH 7.5, 10 mM KOAc, 1 mM Mg(OAc)_2_, 4 mM DTT, and Complete EDTA-free protease inhibitor cocktail (Roche) at an equal volume to the size of the pelleted cells. After 5 min of incubation on ice, cells were transferred into a pre-cooled Dounce homogenizer and lysed by 20-25 strokes. The lysate was centrifuged at 10,000*g for 10 min at 4 °C, and the supernatant transferred to a fresh tube. Aliquots were flash frozen in liquid N_2_ and stored at −80°C.

#### In vitro translation assays

In vitro translation reactions were performed as previously described (32–34), with modifications relating to the Nsp1 addition. HEK293F-cell lysate was pre-incubated with increasing concentrations (from 0 to 3 μM final concentration) of recombinant Nsp1 (wild-type and variants) for 15 min on ice. Translation buffer (20 mM HEPES-KOH, pH 7.6, 1 mM DTT, 0.5 mM spermidine-HCl, 1 mM Mg(OAc)_2_, 8 mM creatine phosphate, 1 mM ATP, 0.2 mM GTP, 150 mM KOAc, 25 μM of each amino acid, and 2 units of human placental ribonuclease inhibitor (Fermentas) was then added followed by 1 μl of the reporter mRNA (0.5 pmol/μl) to give a total reaction volume of 13 μl. Final RNA concentration in the reaction mixture was kept at 38 nM. Translation reactions were incubated at 30°C for three hours, samples were flash frozen in liquid N_2_ and kept at −80°C. Luciferase assays were performed using the Steady-Glo Luciferase Assay kit (Promega) following the manufacturer’s protocol. Luminescence was measured using the M200 finite series microplate reader (TECAN). Samples for all concentrations of each protein were prepared in triplicates and measured.

#### Thermal shift assay

All proteins were used at a final concentration of 1 mg/ml in a total volume of 20 μl. SYPRO Orange (Invitrogen S6651) was used at a final concentration of 5x for all the samples. Experiments were carried out in 20 mM HEPES-KOH, pH 7.51, 200 mM KOAc, 2 mM Mg (OAc)_2_, 0.1 mM THP, and 2.5% glycerol. Each sample was prepared in triplicates. Samples were dispensed into FrameStar 96 well PCR plate (4titudE 4ti-07 10/C) sealed afterward with PCR optical Seal (4titudE). Thermal scanning (10 to 95°C at 1.5 °C/min) was performed using a real-time PCR instrument C1000 Touch Thermal Cycler (CFX96 from Bio-Rad) and fluorescence intensity was measured after every 10 seconds. According to the described protocol, raw data were truncated in Microsoft Excel to remove post-peak quenching (35). A non-linear fitting of the truncated dataset to a Boltzmann Sigmoidal equation was performed to obtain the melting temperature (T_m_) using Prism 9 (GraphPad Software).

#### Structure predictions

All structure predictions were performed using AlphaFold 2 (30) as implemented in ColabFold (https://colab.research.google.com/github/sokrypton/ColabFold/blob/main/AlphaFold2.ipynb) with default settings and allowing the structures to relax using the built-in amber (36) functionality.

#### Molecular Graphics and visualisation

Cartoon representations of Nsp1 structures were generated using ChimeraX (37). Schematic representations of structure topology were drawn in TopDraw (38).

## Acknowledgements

P.K. was supported by a Kempe Foundations postdoctoral fellowship, E.S. was supported by a Umeå University Excellence by Choice (EC) postdoctoral fellowship. L.A.C. was supported by the Human Frontier Science Program (Career Development Award CDA00047/2017-C) and the Knut and Alice Wallenberg Foundation (through the Wallenberg Centre for Molecular Medicine Umeå). V.H. was supported by grants from Cancerfonden (20 0872 Pj) and the Swedish Research council (2017-03783). The project was supported through an exploratory grant for COVID-19-related research, Medical faculty at Umeå University, 2020 to L.A.C. and V.H..

