## Supplementary Information for "Clinically observed deletions in SARS-CoV-2 Nsp1 affect protein stability and its ability to inhibit translation"

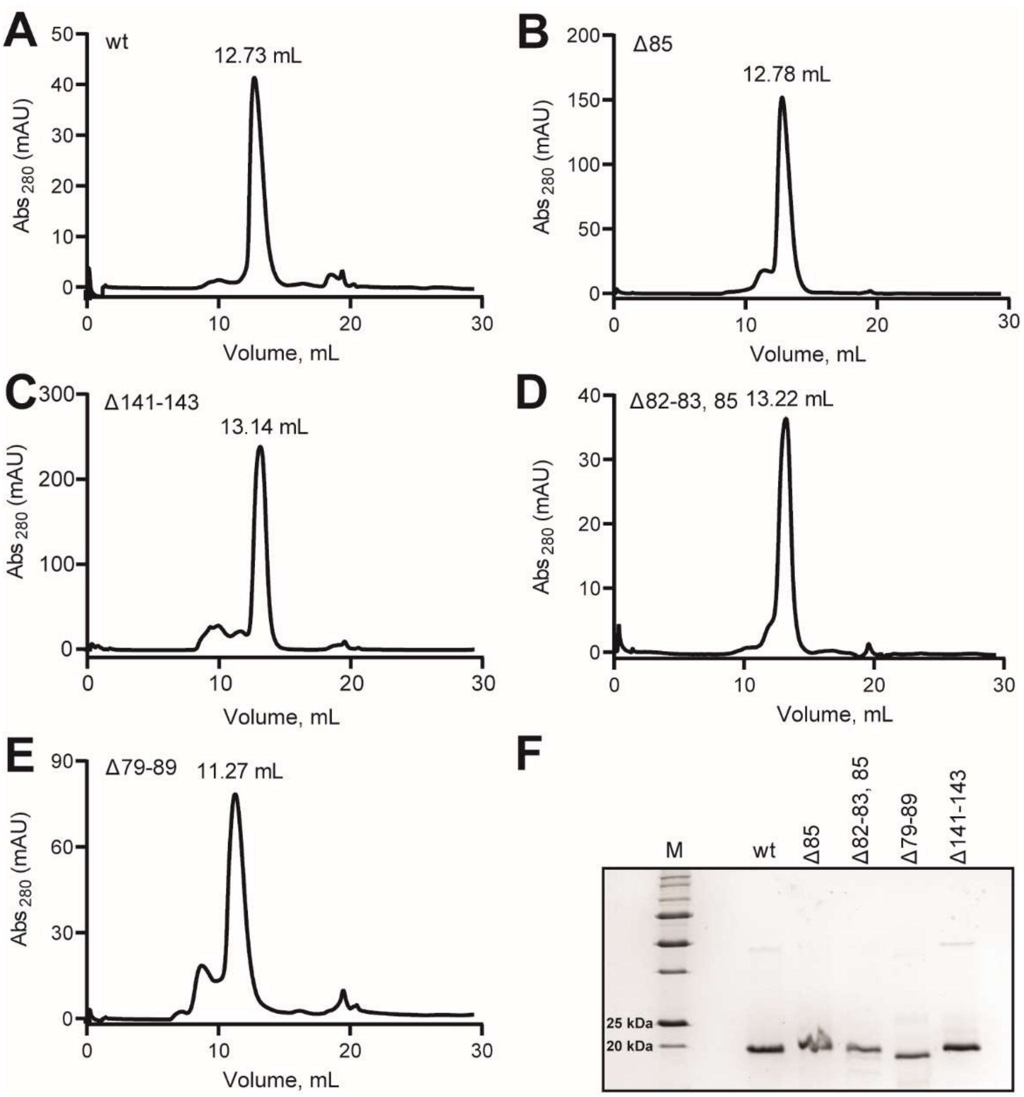


**Figure S1: Purification of wild type and deletion variants of SARS-CoV-2 Nsp1. (A-E)** Chromatograms (280 nm absorption) of Nsp1 and its deletion variants eluting from a Superdex 75 increase 10/300 size-exclusion column. Each chromatogram has the volume corresponding to the center of the main peak noted. The $A_{260}/A_{280}$ ratio was similar (~0.6) in the main peak of all constructs. **(F)** 1.5 µg of each protein sample was analyzed on a 15% SDS-PAGE and stained with Coomassie blue.

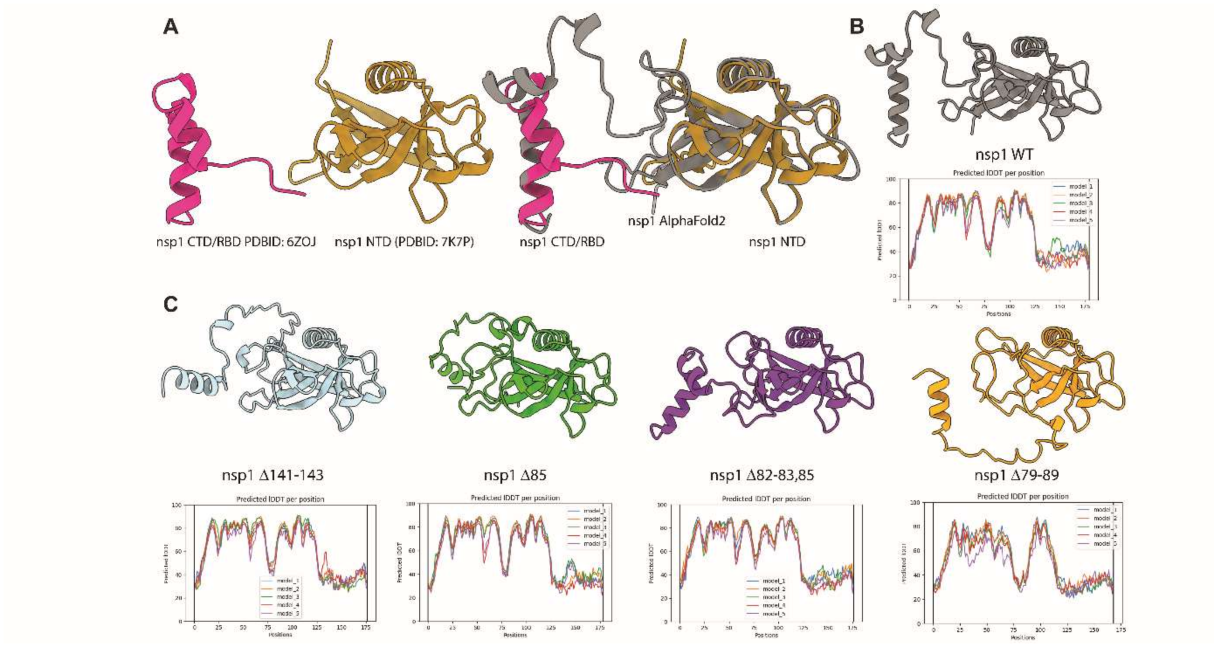


**Figure S2: AlphaFold 2-predicted Nsp1 structures.** **(A)** (*left*) Structures of the experimentally determined N- (residues 10-126, PDBID: 7K7P, (1)) and C-terminal (residues 148-180 PDBID: 6ZOJ, (2)) domains of Nsp1. Structures are oriented as the predicted full-length structure of wt Nsp1 in (B). (*right*) Superposition of the predicted wt Nsp1 structure with the experimentally determined structures as in (A). **(B)** Predicted structure of full length wt Nsp1 shown in cartoon representation (grey) and the corresponding per-residue confidence score (pLDDT) trace for all five prediction runs. **(C)** Cartoon representations of the predicted structures of Δ141-143 (light blue), Δ85 (green), Δ82-83,85 (purple) and Δ79-89 (orange) Nsp1 and their respective per-residue confidence score (pLDDT) trace for each of their predictions.
